# Convergent metatarsal fusion in jerboas and chickens is mediated by similarities and differences in the patterns of osteoblast and osteoclast activities

**DOI:** 10.1101/688036

**Authors:** Haydee L. Gutierrez, Rio Tsutsumi, Talia Y. Moore, Kimberly L. Cooper

## Abstract

The extraordinary malleability of the vertebrate limb supports a variety of locomotor functions including running and leaping in cursorial and saltatorial species. In many of these animals, the metatarsals and/or metacarpals are disproportionately elongated to increase stride length and fused into a single larger element, likely to resist fracture due to increased ground reaction forces. Despite the fact that metapodial fusion evolved convergently in modern birds, ungulates, and jerboas, the developmental basis has only been explored in chickens, which diverged from the mammalian lineage approximately 300 million years ago. Here, we use the lesser Egyptian jerboa, *Jaculus jaculus*, to understand the cellular processes that unite three distinct metatarsal elements into a single cannon bone in a mammal, and we revisit the developing chicken to assess similarities and differences in the localization of osteoblast and osteoclast activities. In both species, adjacent metatarsals align along flat surfaces, osteoblasts cross the periosteal membrane to unite the three elements in a single circumference, and osteoclasts resorb bone at the interfaces leaving a single marrow cavity. However, although spatial and temporal partitioning of osteoblast and osteoclast activities reshape three bones into one in both species, the localization of osteoclasts is distinct. While osteoclasts are uniformly distributed throughout the endosteum of chicken metatarsals, these catabolic cells are highly localized to resorb bone at the interfaces of neighboring jerboa metatarsals. Each species therefore provides an opportunity to better understand the mechanisms that partition osteoblasts and osteoclasts to alter the shape of bone during development and evolution.

## 1. Introduction

A diversity of limb skeletal forms support the weight of the body and allow a range of locomotor activities in vertebrate animals. In species that run or hop at high speeds, the metatarsals and/or metacarpals are often disproportionately elongated to increase stride length. However, simply lengthening a rod-like structure is accompanied by an increase in the likelihood of failure, or fracture, due to an increased bending moment with respect to a point load (e.g. an animal’s foot striking the ground with metatarsals at an angle that is less than perpendicular to the ground). Vertebrate limbs have adopted a variety of bone densities, cortical thicknesses, and curvatures to compensate for increased length in order to maintain an optimal safety factor that balances stiffness while minimizing weight (Brassey et al., 2013). In cases of extreme elongation in multiple lineages that include the birds, jerboas, and artiodactyls (e.g. camels and deer), metatarsals have fused to transform multiple thin rod-like elements into a single ‘cannon bone’ with a larger outer diameter (Clifford, 2010; Moore et al., 2015; Mayr, 2016). Even if comprised of the same material properties with the same cortical thickness, the distribution of mass further from the center of the cross section (neutral bending axis) would increase the second moment of area and the stiffness of the metatarsus (Koch, 1917). Therefore, a fused cannon bone would be expected to withstand higher bending forces than unfused metatarsals.

In vertebrates, the limb skeletal primordia emerge from within limb bud mesenchyme as small cartilage condensations. Each cartilage forms a template that is ultimately replaced by bone through the process of endochondral ossification. Chondrocytes in the center of the cartilage enlarge to become hypertrophic chondrocytes that are then replaced by osteoblasts to form the bony diaphysis, or shaft, with cartilage growth zones at each end of the skeletal element (Kronenberg, 2003). Together, osteoblasts and osteoclasts contribute to normal bone homeostasis, circumferential growth, and reshaping; osteoblasts have an anabolic activity and deposit new bone whereas osteoclasts are catabolic cells that resorb mineralized bone matrix (Hadjidakis and Androulakis, 2006).

While longitudinal growth of each limb bone is driven by the activity of growth cartilages, circumferential growth is driven by the activity of osteoblasts and osteoclasts that line the inner (endosteal) and outer (periosteal) surfaces of cortical bone. Radial growth is one type of circumferential growth of the long bone wherein periosteal mineral deposition is equally countered by resorption on the endosteal surface resulting in a relatively consistent cortical thickness. In contrast, preferential periosteal growth results in an increased cross-sectional thickness, while non-uniform circumferential growth alters the cross-sectional shape and/or produces a linear curvature (Bateman, 1954; Allen and Burr, 2014).

Post-ossification fusion to unite the metatarsals into a single bone evolved independently in multiple vertebrate lineages, including bipedal hopping jerboas, the ancestor of modern birds, and artiodactyls that have additionally fused the metacarpals. Both in mammals and in birds, the metatarsals first develop as individual cylindrical bones that later align and fuse the cortices into one bone that shares a single marrow cavity (Namba et al., 2010; Cooper et al., 2014; Lopez-Rios et al., 2014). Despite its frequent occurrence throughout vertebrate evolution, the developmental basis of bone fusion has been characterized only during fetal chicken development. However, the deep divergence of the synapsid (mammal) and saurapsid (dinosaurs including bird-like ancestors) lineages more than 300 million years ago means there is little reason to assume that the process of bone fusion is similar in birds and in mammals.

Here, we use the bipedal three-toed jerboa as a model to investigate the cellular mechanisms of bone fusion in a mammal. There are 33 species of jerboas including two genera of pygmy jerboas with metatarsals that do not fuse (*Salpingotus* and *Cardiocranius*), one species in which the three central metatarsals are partially fused (*Euchoreutes naso*), and a majority of jerboas with the three central metatarsals that fully fuse into a single cannon bone (Moore et al., 2015). The species that we study to understand the evolution of limb development, *Jaculus jaculus*, does not complete formation of the first and fifth digits during early embryonic development (Cooper et al., 2014). At birth, the remaining metatarsals of digits II, III, and IV are separate cylindrical bones. Soon after birth, while the metatarsals continue to rapidly elongate and grow radially, they fuse to form a single bone in the adult that trifurcates distally and articulates with each of the three proximal phalanges.

We describe the developmental process of metatarsal fusion in the jerboa in three steps: 1) alignment of the three metatarsals to form a transverse arch, 2) formation of mineralized bridges across the periosteum at the interface of adjacent bones, and 3) catabolism of all bone at the interfaces by localized osteoclast activity that unifies a single marrow cavity. To compare and contrast these cellular processes in jerboa with metatarsal fusion in birds (Namba et al., 2010), we also re-assessed chicken metatarsal fusion focusing on bone formation and resorption activities.

In both species, the interfaces of adjacent bone are closely abutted, and mineralized bridges form across the periosteal membrane to unite all three in a single cortical circumference. Bone is removed from the center to unify three distinct cavities into a single marrow space. There are, however, differences in the localization of anabolic (osteoblast) and catabolic (osteoclast) activities. During later phases of bone fusion in the jerboa, osteoblasts remain active at the interfaces of adjacent metatarsals, while osteoclasts are highly localized to degrade bone at the interfaces. In contrast, osteoclasts in the chicken are uniformly distributed around the circumference of all three skeletal elements. Our comparative anatomy and histology explain the structural reorganization of the distal limb to enable bipedal locomotion in the jerboas and in birds and establish the jerboa as a potential model to understand the uncoupling of anabolic and catabolic activities that shape the vertebrate skeleton during development and evolution.

## 2. Materials and Methods

### 2.1 Animals

Jerboas of the species *Jaculus jaculus* were housed and maintained at the University of California San Diego (UCSD) as previously described (Jordan et al., 2011) and in full compliance with the Institutional Animal Care and Use Committee (IACUC). Fertilized chicken eggs (*Gallus gallus*) were purchased from AA Lab Eggs, Inc. and incubated in a humidified rocking cabinet incubator at 37°C until P20 (hatching). Postnatal day 2 (P2) chicken feet were purchased from Charles River Laboratories.

### 2.1 Tissue Processing

Jerboas were anesthetized with 625 mg/kg ketamine 12.5 mg/kg xylazine by intraperitoneal injection and perfused with 20 ml of PBS followed by 20 ml of 4% PFA in PBS at room temperature. Metatarsals were then dissected and placed in 20% sucrose + 4% PFA in PBS solution for 1.5 hours rocking at 4°C. Metatarsals were then embedded in SCEM media (Section-Lab Cat C-EM001) and frozen in a cryomold submerged in an isopentane and dry ice bath. Chicken embryos were collected and stages confirmed according to (Bellairs and Osmond, 2005). Metatarsals were dissected and fixed overnight in 4% PFA in PBS solution then placed in 30% sucrose in PBS solution rocking overnight at 4°C. Metatarsals were then embedded in SCEM media (Section-Lab Cat C-EM001) and frozen in a cryomold suspended in an isopentane and dry ice bath.

Frozen specimen blocks were sectioned on a Leica Cryostat CM1950 using the CryoJane Tape-Transfer System. Adhesive Tape Windows (Leica Cat 39475214) were used to pick up the sections, which were then placed on slides coated with Solution B (Leica Cat 39475271). Slides were then exposed to two pulses, eight milliseconds each and at 360 nm of UV light, and tape was removed.

### 2.2 Tartrate resistant acid phosphatase (TRAP) staining

All chemicals used to stain for tartrate resistant acid phosphatase activity (Hadler et al., 2008) were purchased from Sigma. TRAP Basic Incubation Medium is comprised of 9.2 g sodium acetate anhydrous, 11.4 g L-(+) tartaric acid (Cat 228729), 950 ml distilled water, and 2.8 ml glacial acetic acid (Cat 695092) adjusted to pH 4.7-5.0 with glacial acetic acid or 5 M sodium hydroxide. Naphthol AS-BI phosphate substrate (Cat AC415310010) is dissolved at 20 mg per ml in ethylene glycol monoethyl ether (Cat 128082). Sections were stained for TRAP activity by placing slides in prewarmed TRAP staining solution mix [200 ml TRAP Basic Incubation Medium, 120 mg Fast Red Violet LB Salt (Cat F3381), and 1 ml Naphthol AS-BI phosphate substrate solution] at 37°C for 25 minutes. Slides were then rinsed in distilled water, counterstained with 0.02% Fast Green (Cat F7252) for 90 seconds, and rinsed in distilled water. After dehydration through a graded series of EtOH, slides were cleared in Xylenes and mounted in Permount media (Cat SP15-100) under glass coverslips.

### 2.3 Von Kossa staining

One percent (1%) Aqueous Silver Nitrate Solution was made by adding 1g of Silver Nitrate (Sigma Cat 56506-256) to 100 ml of distilled water. 5% Sodium Thiosulfate solution was made by adding 5g of Sodium Thiosulfate to 100 ml of distilled water. 0.1% Nuclear Fast Red Solution was made by adding 0.1g Nuclear Fast Red (TCI Cat N0305) and 5g Aluminum Sulfate (Spectrum Cat A1114) to 100 ml distilled water and then boiled, cooled, and filtered. A grain of thymol was added as a preservative. Sections were then warmed at 37°C for 15 min, rinsed in PBS 2 × 5 min, then rinsed in distilled water for 1 min. Sections were then incubated in 1% Silver Nitrate Solution in a glass staining dish under ultraviolet light (Spectronics Spectrolinker XL-1000 UV Crosslinker) for 25 minutes. Slides were rinsed in distilled water for 1 minute, then placed in 5% Sodium Thiosulfate, then rinsed in distilled water for 1 minute. Slides were then counterstained in 0.1% Nuclear Fast Red solution for 5 minutes and rinsed in distilled water. Slides were dehydrated 3 minutes in 95% EtOH and 2 × 3 min in 100% EtOH. Slides were then cleared in Xylenes and mounted using Permount media (Fisher Cat SP15-100).

### 2.4 Immunohistochemistry

Slides were washed in PBS 2 × 5 min. Antigen retrieval was then performed by microwaving slides in 1x Epitope Unmasking Buffer (Bioworld cat 21760005-1) for 2 min at a temperature just below 100°C in a glass staining dish. Covered staining dish was then wrapped with a blue underpad and foil for 10 minutes at room temperature, placed at 4°C for 20 minutes, then washed in PBS 2 × 5 min. Slides were placed in blocking solution consisting of 5% heat inactivated goat serum, 0.1% Triton (VWR cat 100504-970), 0.02% SDS for 1 hr at room temperature. Slides were then incubated with primary antibodies anti-Pro-collagen I (DSHB cat SP1.D8) 1:20, anti-Collagen XIVA1(Novus NBP1-86877) 1:250, or anti-Periostin 1:100 (Abcam cat ab14041) diluted in block solution overnight at 4°C. Slides were washed in PBS + 0.1% Triton 3 × 10 min. Slides were then incubated in the dark in Alexa Fluor secondary antibodies goat anti-mouse 488 (Life Tech Cat A21121) 1:500, goat anti-rabbit 594 (Life Tech Cat A11012) 1:500, and 1 µg/ml DAPI (Life Tech Cat D1306) diluted in block solution for 1 hour at room temperature. Slides were then washed in PBS + 0.1% Triton 3 × 10 min and PBS 1 × 5 min then coverslipped with Fluoromount G (Southern biotech Cat OB10001) under glass cover slips. All images were taken on an Olympus BX61 compound microscope.

### 2.5 Computed Tomography

Skeletal specimens for *Cardiocranius paradoxus* (MSB 199763) and *Euchoreutes naso* (MSB 227347) were mounted in floral foam and fluid specimens for *Jaculus jaculus*, obtained from the colony at the Harvard Concord Field Station, were mounted in a plastic container in 95% ethanol for micro-computed tomography (micro-CT) scanning. The specimens were scanned with a SkyScan 1173 micro-CT scanner (Bruker microCT, Kontich, Belgium). Specimens were scanned with 70 kV voltage and 114 µA current for all specimens except for the adult *J. jaculus*, which was scanned at 60 kV and 113 µA. Specimens were scanned at resolutions resulting in 26.29 µm (*Cardiocranius paradoxus*), 22.3 µm (*Euchoreutes naso*), 16.34 µm (adult *J. jaculus*), 35.53 µm (juvenile *J. jaculus*) pixel sizes. The section images were reconstructed with the program NRecon and exported as 3D surface models. We segmented the surface models to isolate the metatarsals and digits and remove the proximal 50% of the metatarsals using MeshLab (Cignoni et al., 2008).

## 3. Results

### 3.1 Metatarsal fusion in bipedal jerboas correlates with increased body size

The earliest diverging extant bipedal jerboas are the pygmy jerboas (subfamily Cardiocraniinae) with adult body weights that range from 6 to 13 g (Shenbrot et al., 2008). These species have metatarsals that closely align in a transverse arch, but each metatarsal remains as a distinct skeletal element in adult animals (Figure 1a). The next most recently derived morphotype is represented here by the long-eared jerboa (subfamily Euchoreutiinae, species *Euchoreutes naso*) with metatarsals that are partially fused; the adult animals, which average about 30 grams in body weight (Stubbe et al., 2007), have a contiguous bone marrow, but remnants of bone at the interfaces of adjacent metatarsals remain as columns that traverse the medullary cavity (Figure 1b). The five-toed (subfamily Allactaginae) and three-toed jerboas (subfamily Dipodinae, including species *Jaculus jaculus* and *Dipus sagitta*) are the most recently divergent; all have fully fused the three central metatarsals into a single cylindrical marrow cavity with no trace of the former interfaces between adjacent bones (Figure 1c, d), other than distally where the three trifurcate into the proximal phalanges.

**FIGURE 1.**
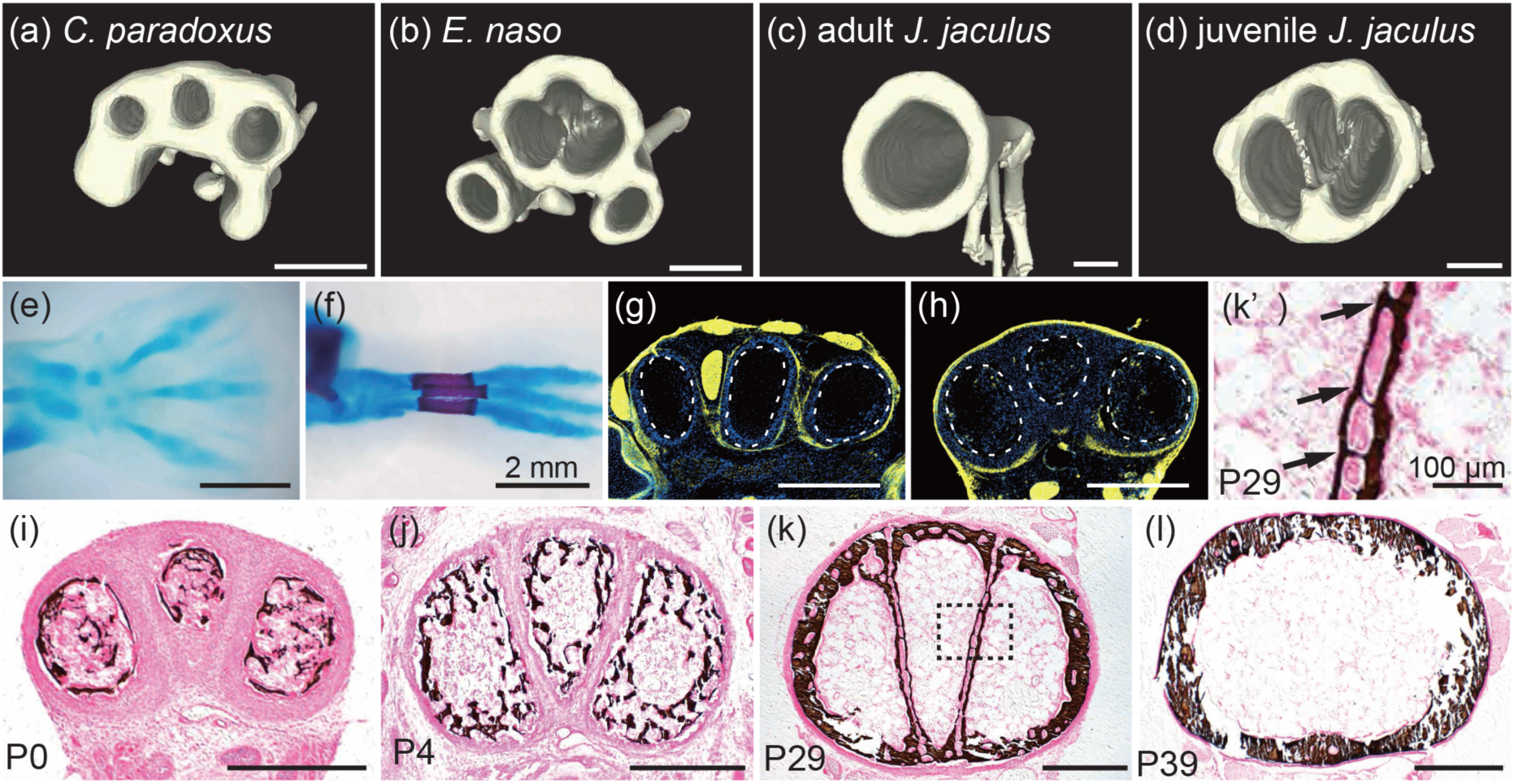
Evolutionary and developmental process of alignment and fusion to unite three metatarsals into a single cannon bone in recently derived jerboas. Micro-computed tomography showing cross-sections at approximately 50% the length of metatarsus III of adult *Cardiocranius paradoxus* (a), *Euchoreutes naso* (b), adult *Jaculus jaculus* (c), and juvenile *Jaculus jaculus* (d). Three-toed jerboa (*Dipus sagitta*) at approximately E13.5 (e) and P0 (f). Immunostaining with anti-Collagen XIV (yellow) and DAPI (blue) on cross-sections of P0 jerboa (*Jaculus jaculus*) hand (g) and foot (h). Dotted lines represent metapodial bones. While there are Collagen XIV positive layers between metacarpals in the hand, a Collagen XIV layer is absent between metatarsals in the foot (n=3 each). (i-l) Temporal dynamics of cross-sectional shapes of metatarsals of the lesser Egyptian jerboa (*Jaculus jaculus*) at P0 (i, n=3), P4 (j, n=3), P29 (k, n=4), and P39 (l, n=3). The samples were stained by Von Kossa to detect mineralized bone in black and counterstained with eosin in pink. (k’) The enlarged view of dotted-square in (k) at P29. Arrows show the bridging of three metatarsals. Each scalebar measures 500 µm unless otherwise indicated in the figure.

Since metatarsals in quadrupedal rodents and in the earliest diverging pygmy jerboas are not fused, the transition from quadrupedalism to bipedalism most likely occurred in species with metatarsals that were also not fused. The unfused metatarsals of pygmy jerboas are disproportionately longer than those of closely related quadrupedal species (Moore et al., 2015), and they would have experienced a greater force per gram of animal while standing stationary by distributing weight among half as many feet. Furthermore, the shift from scurrying to hopping involves the addition of aerial phases, which produce much greater ground reaction forces and bone stresses, even if the same number of legs are used in each gait (McMahon et al., 1987; Rubenson et al., 2004). We therefore hypothesize that the lack of fusion may have constrained the body size to maintain a safety factor that prevents metatarsal fracture. Indeed, increasingly fused metatarsals are accompanied by increases in body size; jerboas with fully fused metatarsals have reached a body size up to 415 grams in *Allactaga major*, two orders of magnitude greater than the pygmy species (Shenbrot et al., 2008). Since ground-reaction force and stress on elongated metatarsals increases with body size and with the addition of aerial phases in locomotion, this pattern suggests that metatarsal fusion may have been a structural adaptation that allowed body size to increase in these bipedal species.

### 3.2 Metatarsal fusion in *Jaculus jaculus* proceeds through a series of events to reshape bone

We describe the developmental process of complete metatarsal fusion in the crown group of jerboas that have fully fused metatarsals using *Dipus sagitta* for late gestational stages and *Jaculus jaculus* for postnatal analyses (Moore et al., 2015; Pisano et al., 2015). In the mid-gestation jerboa embryo, metatarsal condensations of digits II and IV each angle away from the metatarsal of digit III (Figure 1e). By birth, the three metatarsals lie alongside one another with the middle metatarsal in a slightly more dorsal position, and each has a circular shape in cross-section (Figure 1f, i). Within days after birth, the cross-sectional shape of each metatarsal has transformed such that the three collectively comprise a transverse arch (Figure 1j). The third metatarsal has adopted a ‘pie-wedge’ shape in cross-section and serves as the ‘keystone’, while the second and third metatarsals each have a more half-circular shape with a flat surface that lies adjacent to the central metatarsal. The neonatal state of three distinct metatarsals aligned into a transverse arch recapitulates the adult morphology of the earliest diverging pygmy jerboas (Figure 1a, i).

In *J. jaculus*, each of the three metatarsals is maintained as a discrete mineralized element for approximately three to four weeks after birth. The first evidence that fusion has begun is the appearance of mineralized bridges that invade the periosteal space and physically connect adjacent metatarsals (Figure 1k, k’ arrows). Mineralized bridging occurs not only dorsally and ventrally to unite the cortex that will ultimately encircle the unified marrow cavity, but also along the length of adjacent interfaces. Within 5-6 weeks after birth, all mineralized bone at these interfaces has been removed, and a single marrow cavity remains encircled by cortical bone (Figure 1l).

Although the initiation of bridging and of bone catabolism do not occur at precise developmental ages or animal weights, their relative order is consistent and proceeds from proximal to distal over time. Transverse sections through an animal at P29 shows that catabolism of proximal bone is more similar to the midshaft of an older animal while the pattern of mineralized bridges more distally in the same individual is similar to more proximal bone of a younger animal (Supplementary Figure 1).

Since osteoblasts invade the periosteal membrane of adjacent metatarsals to form mineralized bridges, we suspected there may be a difference in the structure of the periosteum itself. In chicken, Collagen XIV, also known as Undulin, marks the outer layer of the periosteum (Bandyopadhyay et al., 2008). In jerboa metacarpals that remain as distinct skeletal elements into adulthood, Collagen XIV expression encircles each skeletal element and is also strongly expressed in tendons and surrounding connective tissue (Figure 1g). In contrast, there is no expression of Collagen XIV between jerboa metatarsals (Figure 1h). Instead, a domain that may be the outer layer of periosteum, or a distinct layer of connective tissue, encircles all three skeletal elements.

### 3.3 Spatial and temporal pattern of anabolic bone deposition by osteoblasts and catabolic bone resorption by osteoclasts in jerboa metatarsals

With an understanding of the reorganization of mineralized bone over time, we next sought to determine if there is a correlated pattern of anabolic osteoblast activity. We performed immunofluorescence with an antibody to detect pro-Collagen I, the intracellular precursor of Collagen I, which is the major protein component of bone and is produced by osteoblasts (Prockop et al., 1979). In newborn jerboas, we saw pro-Collagen I expression in cells that completely encircle each of the three metatarsals in transverse sections (Figure 2a, b). However, by postnatal day 21, expression is discontinuous in the periosteum that lines the interface of adjacent metatarsals, though pro-Collagen I is present along the endosteal surface of bone at the interfaces (Figure 2c, c’, c” arrow and arrowhead, respectively). We frequently observe no pro-Collagen I expression in the dorsal-most aspect of periosteum that lies between adjacent metatarsals (Figure 2c’, arrow).

**FIGURE 2.**
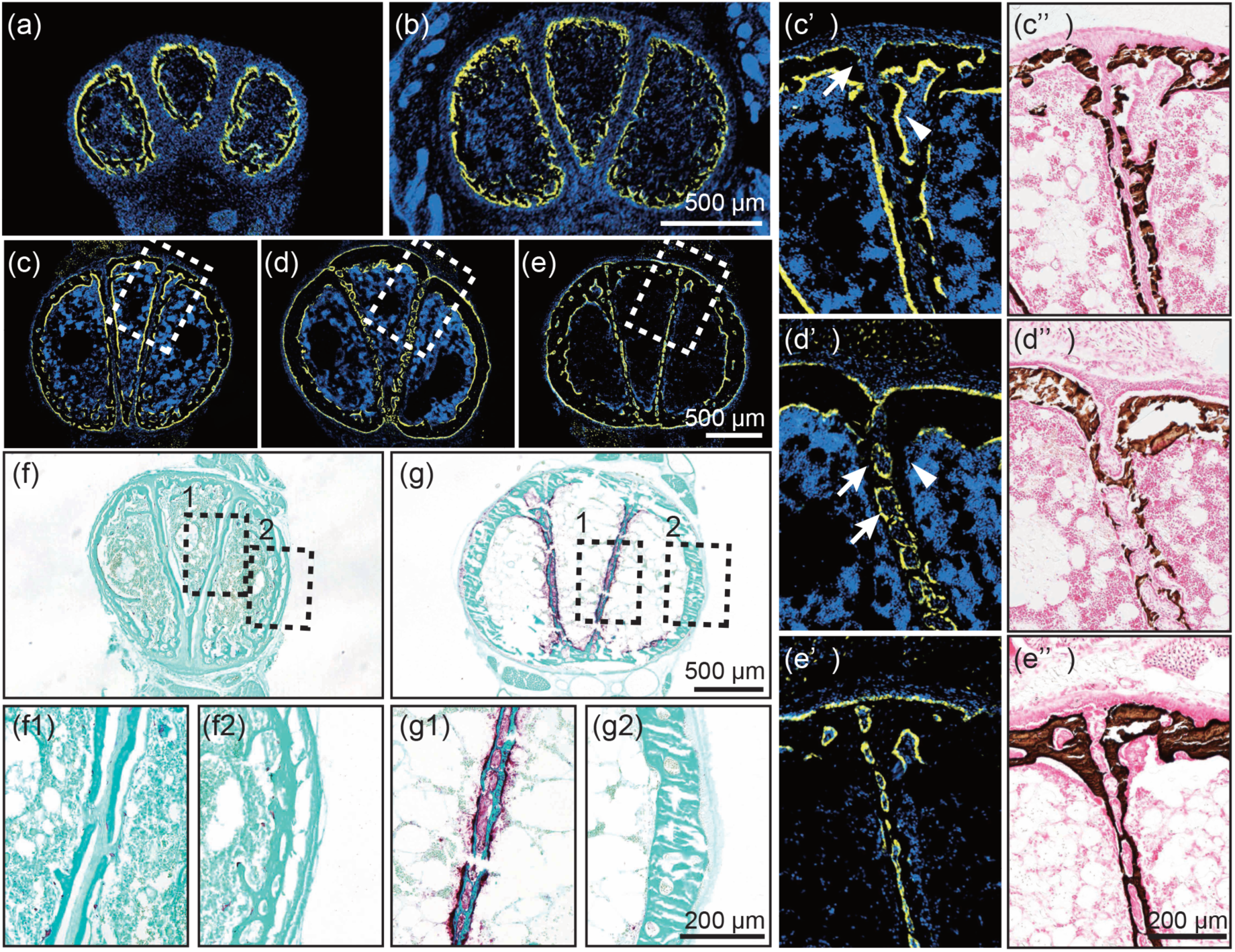
Temporal and spatial localization of osteoblasts and osteoclasts in metatarsals of jerboa (*Jaculus jaculus*). Immunostaining using anti-pro-Collagen I (yellow) and DAPI (blue) of metatarsals in the lesser Egyptian jerboa at P0 (a, n=5), P4 (b, n=4), P21 (c, n=5), P26 (d, n=2), and P29 (e, n=2). The enlarged views of the dotted rectangles in (c), (d), and (e) are shown in (c’), (d’), and (e’). Von Kossa staining of corresponding serial sections are shown in (c”), (d”), and (e’’) respectively. Arrow in (c’) denotes absence of the pro-Collagen I in the dorsal-most periosteum, and arrowhead shows expression in the endosteal layer. Arrows in (d’) show the expression of pro-Collagen I in the periosteum between metatarsals, and arrowhead shows absence of expression in the endosteal layer. TRAP staining of metatarsals at P14 (f, n=4) and P29 (g, n=4) detects osteoclast activity in maroon counterstained with Fast Green. The enlarged views of the dotted rectangles are shown in (f1 and f2) and (g1 and g2), respectively. Scalebars denote measurements for groups of related panels.

The pattern of pro-Collagen I expression along interfaces between adjacent bones during the fourth week is somewhat surprising in light of the apparent downregulation of pro-Collagen I expression in the periosteum by around three weeks after birth. At this later time point, expression becomes absent from the endosteum, instead lines the periosteum, and in fact crosses the periosteum (Figure 2d, d’ arrowhead and arrows, respectively). This pattern of expression at four weeks is consistent with our observation that mineralized bridges cross the periosteum between adjacent metatarsals, indeed precedes the pattern of mineralization, and appears to encircle segments of the periosteum (Figure 2d”). To determine if cells within these loops of pro-Collagen I expressing cells retain periosteal identity, we also detected the expression of Periostin, an extracellular matrix protein expressed in connective tissues that include the periosteum. We find Periostin expression that is encircled by osteoblasts that line the mineralized bridges in the space between adjacent metatarsals as well as strong and contiguous Periostin expression surrounding all three metatarsals (Supplementary Figure 2a, b).

Osteoclast activity can be detected by enzyme chemodetection of tartrate resistant acid phosphatase (TRAP). From soon after birth until near the end of the fourth postnatal week, there is very little osteoclast activity at the mid-diaphysis (Figure 2f, f1, f2). Beginning at around four weeks after birth, we observed high osteoclast activity associated with bone that lies at the interfaces of the three metatarsals (Figure 2g, g1). In contrast, we observed very little TRAP activity associated with the outer cortical bone that encircles what is becoming a single marrow cavity (Figure 2g2). At this stage, there is still expression of pro-Collagen I at the interface of the three metatarsals (Figure 2e, e’, e”). Together, this suggests that osteoclast differentiation or recruitment is localized to restrict bone resorption to the interfaces of adjacent bone within what will become a single marrow cavity.

### 3.4 Osteoblast and osteoclast activities during fusion of the tarsometatarsus in chickens

We showed that metatarsal fusion in jerboas occurs by locally altering the patterns of bone deposition and degradation through temporal and spatial differences in the anabolic activity of osteoblasts and the catabolic activity of osteoclasts. Bone fusion also occurs during late fetal development in all modern birds to unite the metatarsals of hindlimb digits II-IV into a single ‘tarsometatarsus’ that also includes some of the tarsal, or ankle, elements. Considering the phenotypic convergence of metatarsal fusion in these two clades that diverged more than 300 million years ago, we next asked whether the independent evolution of metatarsal fusion occurred by similar spatial partitioning of osteoblast and/or osteoclast activities.

Although the morphological process of tarsometatarsal fusion in chicken was previously well-described (Namba et al., 2010), we re-evaluated the shape transformation over time to set the stage for our investigation of osteoblast and osteoclast activities. As in this prior work, Von Kossa staining of mineralized bone in transverse sections through the chicken foot at embryonic day 13 (E13) revealed three distinct metatarsals each with a circular shape in cross-section through the midpoint of the diaphysis, the position where metatarsal fusion initiates in chicken (Figure 3a). By E15, the metatarsals of digits II and IV have adopted a half-circular shape, and the metatarsal of digit III has become square at this position. Although the three metatarsals of the chicken foot lie in a plane alongside one another and do not form a transverse arch as in the jerboa, neighboring metatarsals of the chicken and jerboa do lie adjacent to one another along flat interfaces (Figure 3b).

**FIGURE 3.**
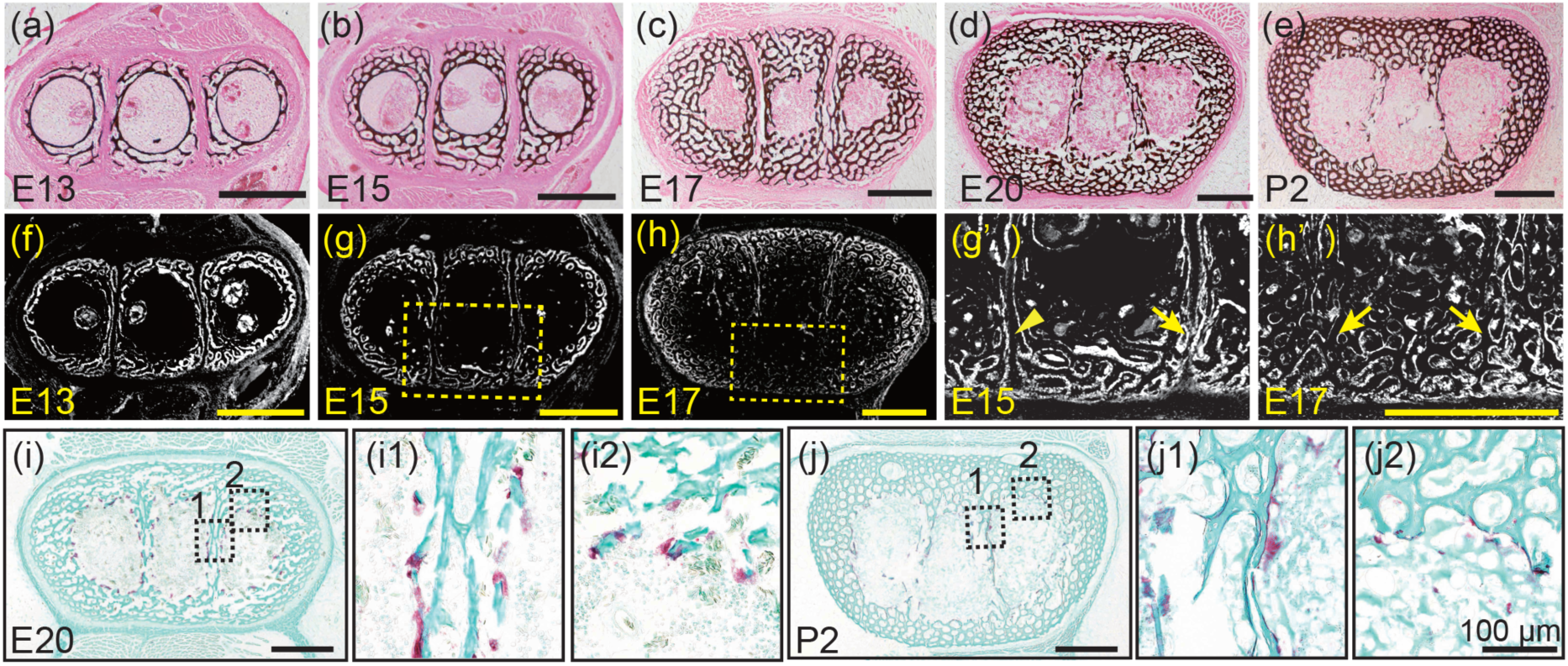
Mineralized bone fusion and temporal and spatial localization of osteoblasts and osteoclasts in metatarsals of chicken (*Gallus gallus*). (a-e) Von Kossa staining of mineralized bone at E13 (a, n=4), E15 (b, n=4), E17 (c, n=4), E20 (d, n=4), and P2 (e, n=2). (f-h’) pro-Collagen I immunostaining (white) in serial sections at E13 (f, n=4), E15 (g, n=4), E17 (h, n=4). The enlarged views of the dotted rectangles in (g) and (h) are shown in (g’) and (h’), respectively. Arrowhead in (g’) indicates a gap between Pro-Collagen I positive layers. Arrows in (g’) and (h’) show bridges of pro-Collagen I. (i-j’) TRAP staining of tarsometatarsals in chicken at E20 (i, n=4) and P2 (j, n=2). The enlarged view of squares at the interface (1) and circumferential endosteum (2) in (i) and (j) are shown in adjacent panel insets. Scalebars measure 500 µm for panels a-j, g’ and h’. Scalebars measure 100 µm for insets i1, i2, j1, and j2.

Chicken metatarsal cortical bone is more trabeculated, or spongy, and less compact than in jerboa. As described by Namba and colleagues, the three metatarsals of the chicken become connected by trabeculae that form along the dorsal and ventral interfaces of embryos from day 17 to day 20 (Figure 3c, d). These trabeculae appear similar to the mineralized bridges that we observe crossing the periosteal membrane between adjacent jerboa metatarsals. Finally, from E20 to P2, mineralization at the interface of neighboring chicken metatarsals is removed leaving a single marrow cavity as in the jerboa (Figure 3d, e).

Also as in the jerboa, the expression pattern of pro-Collagen I appears to predict the structure of mineralized bone in the chicken tarsometatarsus. At E13, expression of pro-Collagen I in tissues surrounding the central metatarsal, which is more circular in its mineralized structure, is similar to the square shape of this skeletal element at E15 (Figure 3b, f). Although mineralized trabeculae do not bridge the space between adjacent metatarsals until about E17, pro-Collagen I expression starts to cross the periosteum by about E15 (Figure 3g, g’, arrow). At E17, pro-Collagen I expression fully crosses the periosteum of adjacent metatarsals (Figure 3h, h’ arrows). At E20, pro-Collagen I expression surrounds the three metatarsals collectively, as trabeculated bone is deposited around the circumference, and we sometimes observed downregulation of pro-Collagen I on the surface of bone that remains at the interfaces (Supplemental Figure 3).

We next assessed whether osteoclast activity is localized to specifically resorb bone at the interfaces of adjacent metatarsals in chickens as in jerboas. In chickens, osteoclasts remain uniformly distributed around the endosteal circumference throughout the process of metatarsal fusion and are not enriched at the interfaces of adjacent metatarsals, in contrast with jerboas (Figure 3i, 3i1, and 3i2). Even from E20 to post-hatching day 2 in the chicken (P2), when bone is mostly resorbed between adjacent metatarsals, osteoclast activity never appears localized to these interfaces (Figure 3j, 3j1 and 3j2). Therefore, the unification of a single marrow cavity in chickens occurs by uniform endosteal osteoclast activity rather than by the highly localized osteoclast activity that we observed in jerboas.

## 4. Discussion

Here, we show that complete metatarsal fusion in the most derived jerboas, as well as in chickens, is characterized by localized alterations of anabolic osteoblast and catabolic osteoclast activities that change the pattern of mineralization in time and in space. Early in the process of bone fusion, the jerboa and chicken share a similar pattern of osteoblast activity that bridges the periosteum between adjacent metatarsals. The unusual behavior of osteoblasts that violate the periosteal boundary in both species is likely necessary to connect all three metatarsals in a single circumferential cortex. Once this is accomplished, osteoblast activity is sometimes decreased along bone interfaces in the chicken but appears to persist in the jerboa until later stages when bone at the interfaces is resorbed in both species.

In the final stage of metatarsal fusion in jerboas, a high density of osteoclast activity lines bone at the interface of adjacent metatarsals, and very few osteoclasts are associated with the endosteal surface that encircles what will be a single marrow cavity. It therefore appears that removal of bone from what will become a single marrow cavity is spatially localized. This contrasts with the chicken, in which osteoclasts appear uniformly distributed along the endosteal surface of each skeletal element. If bone is indeed resorbed uniformly from the endosteal surfaces of chicken metatarsals, it is possible that fusion is achieved, at least in part, by the difference in bone thickness in the two species. Cortical bone surrounding each of the three metatarsals is thin in the jerboa compared to the circumference of highly trabeculated bone that encircles the three elements of the chicken. Bone that lies between adjacent chicken metatarsals is also much thinner, perhaps in part due to decreased osteoblast activity prior to fusion. Uniform bone resorption of a cylinder with non-uniform thickness in the chicken would indeed preferentially remove thin bone at the interfaces, also because the interface is lined by osteoclasts flanking endosteal surfaces while the circumferential bone has only a single endosteal surface.

Prior to metatarsal fusion to form a single marrow cavity in either species, the three metatarsals achieve an intimate alignment that closely abuts adjacent surfaces along flat interfaces. It is reasonable to assume that the close association of periosteal surfaces might be necessary to unify the three skeletal elements into a single cortex, but evidence from the jerboa lineage suggests that such a close association is not sufficient to trigger fusion. The very similar shape of a transverse arch that we observe in neonatal jerboas of the species *J. jaculus*, is also present in adults of the pygmy jerboa species *Cardiocranius paradoxus* (Moore et al., 2015). If close alignment along a flat surface were sufficient to allow passive fusion of adjacent cortical bones, the pygmy jerboa should not have three distinct metatarsals.

How is the locally biased pattern of osteoblast and osteoclast activity established to reshape mineralized bone for fusion? In other species, preferential periosteal growth is affected by mechanical stimuli applied to bones such that the shapes of bones are developmentally adapted to withstand load-bearing demands (Epker and Frost, 1966; Levenston et al., 1998; Frost, 2000). In animals that develop without limb muscle or with muscle paralysis, absence of mechanical loading causes bones to retain a simpler columnar shape, and the mechanical resistance of bones is compromised (Sharir et al., 2011). However, the process of bone fusion by mineralized bridge formation across the periosteal boundary and localization of osteoclast activity occurs prior to the time when juvenile jerboas adopt a quadrupedal gait at about four weeks after birth. Juvenile jerboas do not become fully bipedal until after metatarsal fusion is complete at about five weeks after birth (Eilam and Shefer, 1997). Similarly, tarsometatarsal fusion in fetal chickens is almost complete prior to hatching (Namba et al., 2010). Therefore, bipedal locomotion does not seem to provide a mechanical stimulus that initiates metatarsal fusion, but rather fusion may be due to a developmental program that is a result of natural selection to optimize the metatarsal safety factor.

It is interesting that osteoclast activity is locally upregulated at bone interfaces in the jerboa while osteoblast activity is sometimes locally downregulated at bone interfaces in the chicken. These data suggest that anabolic and catabolic activities can be uncoupled in multiple ways to reshape skeletal elements. In mouse ribs, *Gdf5* expression appears to be controlled by multiple enhancers that partition expression into subdomains along the circumference of the perichondrium (Guenther et al., 2008). Since the developmental process of metatarsal fusion in jerboas and chickens appears to be genetically determined, rather than induced by locomotory activities, these two species may be powerful systems to identify genes and associated cis-regulatory elements that partition the activities of osteoblasts and osteoclasts to define adult bone shape. In addition to providing a basis for understanding the extraordinary diversity of limb bone shapes in vertebrate species, these studies would expand our understanding of two cell types that are critical regulators of bone growth and mineral homeostasis.

## Acknowledgements

We are grateful to Julio Rojano for assistance with cryosectioning. We thank all members of the Cooper laboratory as well as Drs. Michael Perry and James Posakony for thoughtful comments on the manuscript. This work was supported by a Searle Scholar Award from the Kinship Foundation, a Pew Biomedical Scholar Award from the Pew Charitable Trusts, and a Packard Fellowship in Science and Engineering from the David and Lucile Packard Foundation awarded to KLC.

---

**Supplemental Figure 1.**
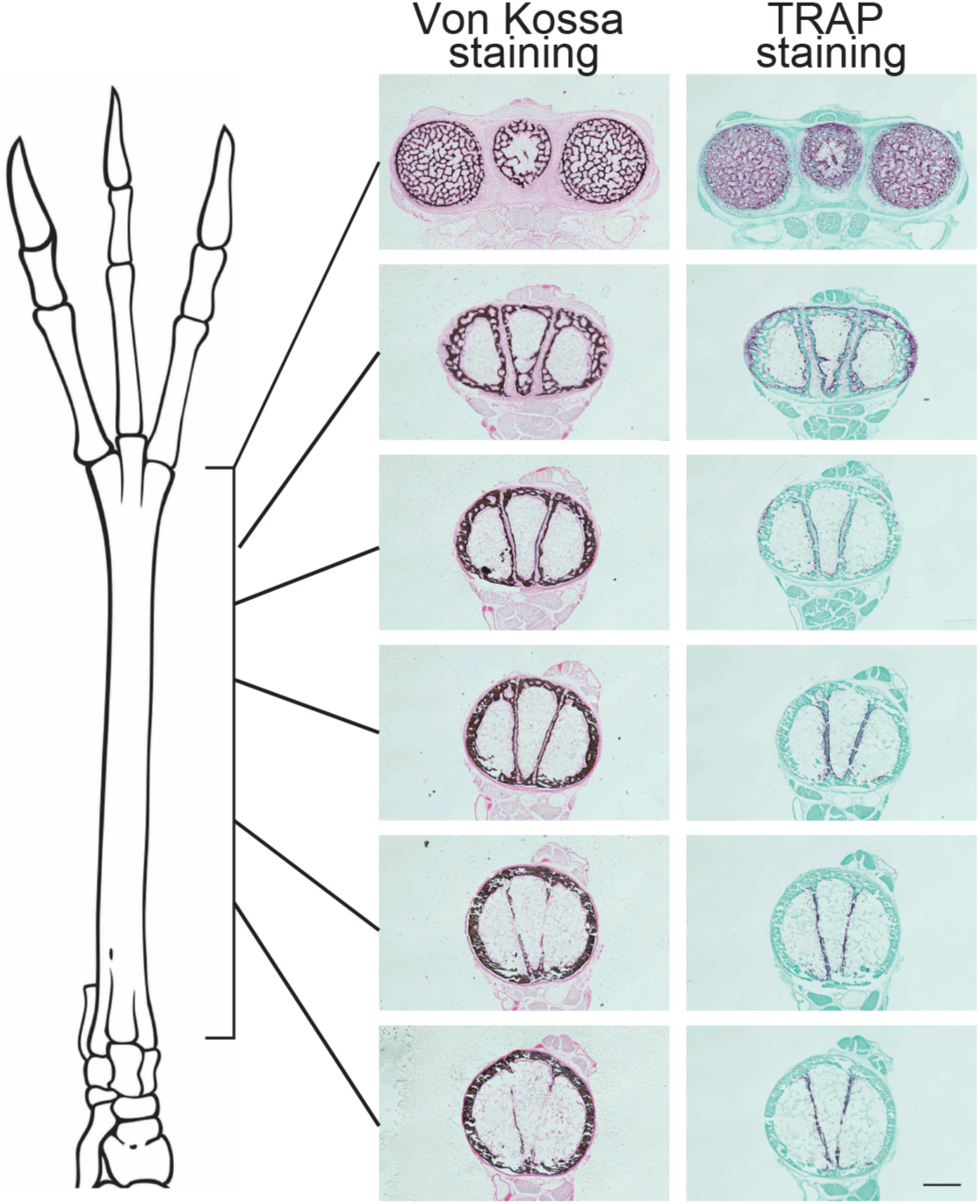
Phases of metatarsal fusion progress from proximal to distal. Von Kossa staining and TRAP staining of transverse sections of P29 jerboa metatarsals from proximal (bottom) to distal (top) positions. n=2. Scalebar = 500 µm.

**Supplemental Figure 2.**
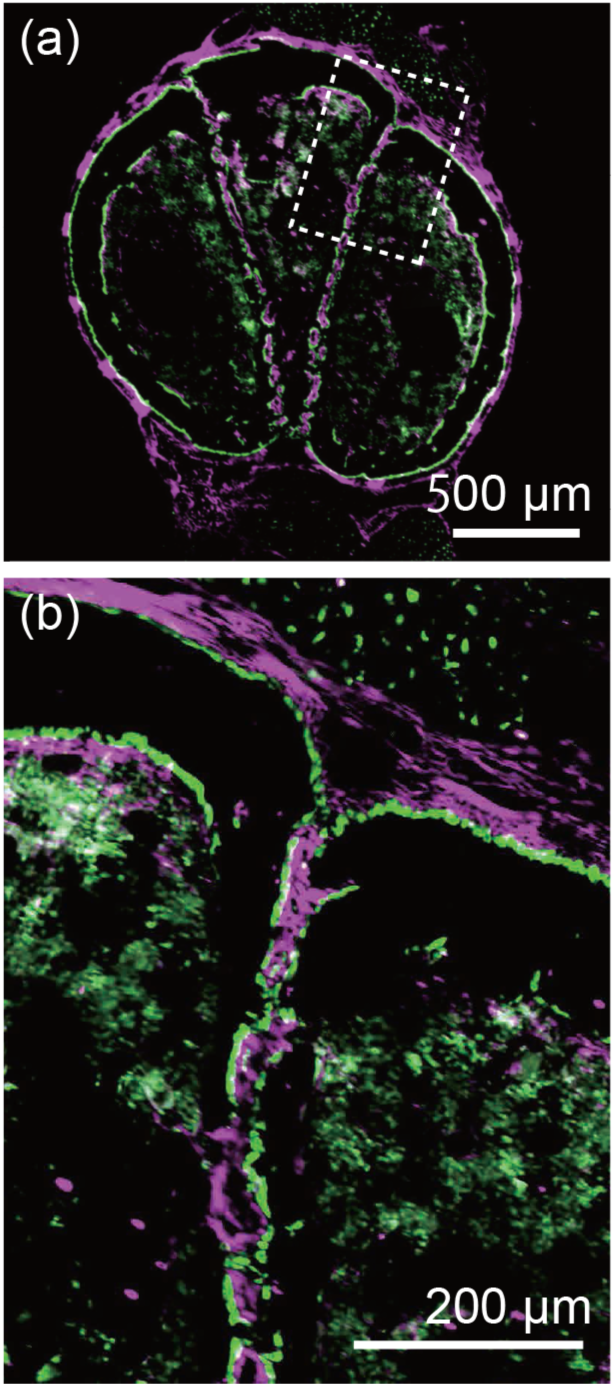
Osteoblasts cross and encircle the periosteum in cross sections. Immunostaining with anti-Periostin (magenta) and Pro-Collagen I (green) of P26 jerboa metatarsals (a). The enlarged view of the dotted rectangle in (a) is shown in (b). n=2. (a) Scalebar = 500 µm. (b) Scalebar = 200 µm.

**Supplemental Figure 3.**
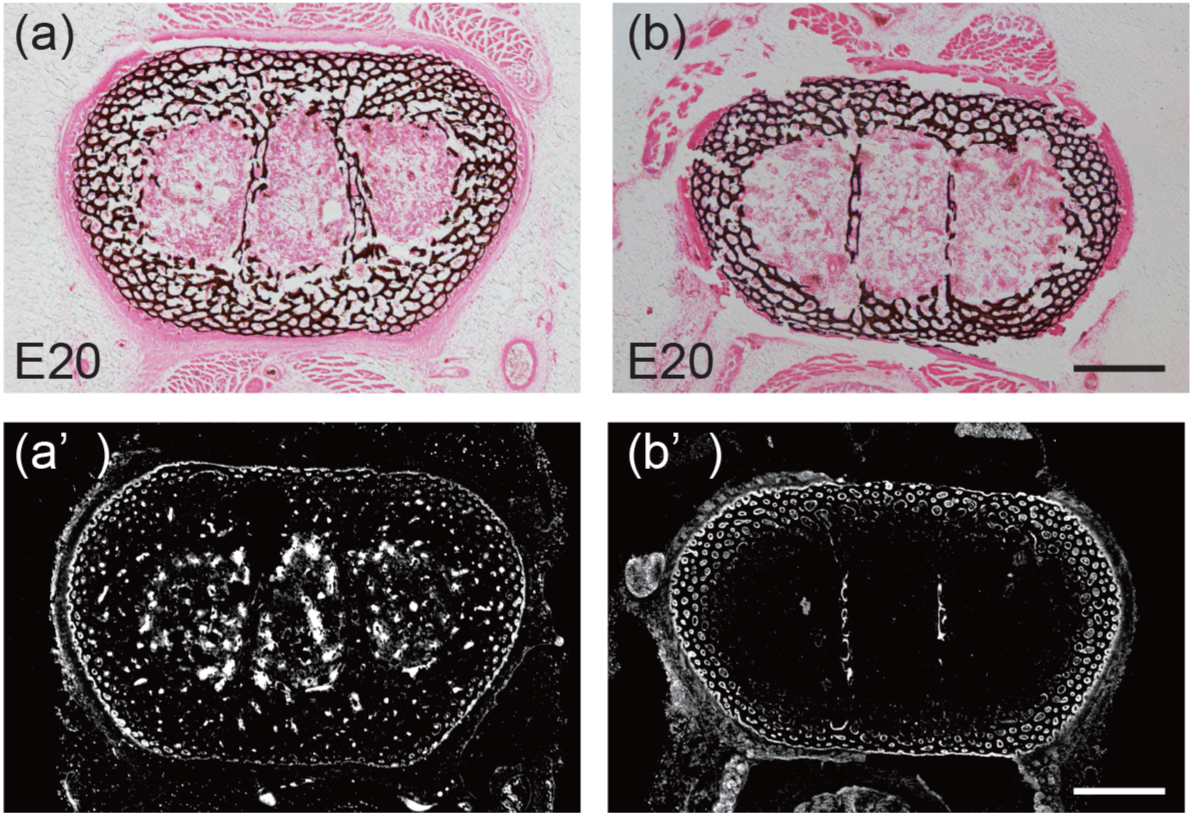
The expression of pro-Collagen I at chicken embryonic day 20 is variable. Von Kossa staining in two individuals of chicken stage E20. Immunostaining with anti-pro-Collagen I (White) on the serial sections of (a) and (b) are shown in (a’) and (b’), respectively. Scalebar = 500 µm.

## References

Allen, M.R., and Burr, D.B. 2014. Chapter 4 - Bone Modeling and Remodeling. In Basic and Applied Bone Biology, D.B. Burr, and M.R. Allen, eds. (San Diego: Academic Press), pp. 75–90.

Bandyopadhyay, A., Kubilus, J.K., Crochiere, M.L., Linsenmayer, T.F., and Tabin, C.J. 2008. Identification of unique molecular subdomains in the perichondrium and periosteum and their role in regulating gene expression in the underlying chondrocytes. Developmental Biology 321: 162–174.

Bateman, N. 1954. Bone growth: a study of the grey-lethal and microphthalmic mutants of the mouse. Journal of Anatomy 88: 212–262.

Brassey, C.A., Kitchener, A.C., Withers, P.J., Manning, P.L., and Sellers, W.I. 2013. The Role of Cross-Sectional Geometry, Curvature, and Limb Posture in Maintaining Equal Safety Factors: A Computed Tomography Study. The Anatomical Record 296: 395–413.

Cignoni, P., Callieri, M., Corsini, M., Dellepiane, M., Ganovelli, F., and Ranzuglia, G. 2008. MeshLab: an Open-Source Mesh Processing Tool (The Eurographics Association).

Clifford, A.B. 2010. The Evolution of the Unguligrade Manus in Artiodactyls. Journal of Vertebrate Paleontology 30: 1827–1839.

Cooper, K.L., Sears, K.E., Uygur, A., Maier, J., Baczkowski, K.-S., Brosnahan, M., Antczak, D., Skidmore, J.A., and Tabin, C.J. 2014. Patterning and post-patterning modes of evolutionary digit loss in mammals. Nature 511: 41–45.

Eilam, D., and Shefer, G. 1997. The developmental order of bipedal locomotion in the jerboa (Jaculus orientalis): pivoting, creeping, quadrupedalism, and bipedalism. Developmental Psychobiology 31: 137–142.

Epker, B.N., and Frost, H.M. 1966. Biomechanical Control of Bone Growth and Development: A Histologic and Tetracycline Study. J Dent Res 45: 364–371.

Frost, H.M. 2000. The Utah paradigm of skeletal physiology: an overview of its insights for bone, cartilage and collagenous tissue organs. J. Bone Miner. Metab. 18: 305–316.

Guenther, C., Pantalena-Filho, L., and Kingsley, D.M. 2008. Shaping Skeletal Growth by Modular Regulatory Elements in the Bmp5 Gene. PLOS Genetics 4: e1000308.

Hadjidakis, D.J., and Androulakis, I.I. 2006. Bone Remodeling. Annals of the New York Academy of Sciences 1092: 385–396.

Koch, J.C. 1917. The laws of bone architecture. American Journal of Anatomy 21: 177–298.

Kronenberg, H.M. 2003. Developmental regulation of the growth plate. Nature 423: 332–336.

Levenston, M.E., Beaupré, G.S., and Carter, D.R. 1998. Loading Mode Interactions in Simulations of Long Bone Cross-Sectional Adaptation. Computer Methods in Biomechanics and Biomedical Engineering 1: 303–319.

Lopez-Rios, J., Duchesne, A., Speziale, D., Andrey, G., Peterson, K.A., Germann, P., Ünal, E., Liu, J., Floriot, S., Barbey, S., et al. 2014. Attenuated sensing of SHH by Ptch1 underlies evolution of bovine limbs. Nature 511: 46–51.

Mayr, G. 2016. Avian Evolution: The Fossil Record of Birds and Its Paleobiological Significance (John Wiley & Sons).

McMahon, T.A., Valiant, G., and Frederick, E.C. 1987. Groucho running. Journal of Applied Physiology 62: 2326–2337.

Moore, T.Y., Organ, C.L., Edwards, S.V., Biewener, A.A., Tabin, C.J., Jenkins, F.A., and Cooper, K.L. 2015. Multiple Phylogenetically Distinct Events Shaped the Evolution of Limb Skeletal Morphologies Associated with Bipedalism in the Jerboas. Current Biology 25: 2785–2794.

Namba, Y., Yamazaki, Y., Yuguchi, M., Kameoka, S., Usami, S., Honda, K., and Isokawa, K. 2010. Development of the Tarsometatarsal Skeleton by the Lateral Fusion of Three Cylindrical Periosteal Bones in the Chick Embryo (Gallus gallus). The Anatomical Record 293: 1527–1535.

Pisano, J., Condamine, F.L., Lebedev, V., Bannikova, A., Quéré, J.-P., Shenbrot, G.I., Pagès, M., and Michaux, J.R. 2015. Out of Himalaya: the impact of past Asian environmental changes on the evolutionary and biogeographical history of Dipodoidea (Rodentia). Journal of Biogeography 42: 856–870.

Prockop, D.J., Kivirikko, K.I., Tuderman, L., and Guzman, N.A. 1979. The Biosynthesis of Collagen and Its Disorders. New England Journal of Medicine 301: 13–23.

Rubenson, J., Heliams, D.B., Lloyd, D.G., and Fournier, P.A. 2004. Gait selection in the ostrich: mechanical and metabolic characteristics of walking and running with and without an aerial phase. Proceedings of the Royal Society of London. Series B: Biological Sciences 271: 1091–1099.

Sharir, A., Stern, T., Rot, C., Shahar, R., and Zelzer, E. 2011. Muscle force regulates bone shaping for optimal load-bearing capacity during embryogenesis. Development 138: 3247–3259.

Shenbrot, G.I., Sokolov, V.E., Heptner, V.G., and Koval’skaya, Yu.M. 2008. Jerboas: Mammals of Russia and Adjacent Regions (Enfield, NH: CRC Press).

Stubbe, A., Stubbe, M., Nyamsuren, B., Samjaa, R., Driechciarz, E., Driechciarz, R., Schonert, A., and Winter, M. 2007. Euchoreutes naso Sclater, 1890 – ein SäugetierEndemit Zentralasiens. Erforschung Biologischer Ressourcen Der Mongolei 471–486.

